# Distribution of malaria parasite-derived phosphatidylcholine in the infected erythrocyte

**DOI:** 10.1101/2023.03.13.532364

**Authors:** Tansy Vallintine, Christiaan van Ooij

**Affiliations:** Department of Infection Biology, Faculty of Infectious Disease, London School of Hygiene & Tropical Medicine, London, United Kingdom

## Abstract

Malaria parasites modify their host erythrocyte in multiple ways, leading to changes in the deformability, adhesiveness and permeability of the host erythrocyte. Most of these changes are mediated by proteins exported from the parasite to the host erythrocyte, where these proteins interact with the host cell cytoskeleton or form complexes in the plasma membrane of the infected erythrocyte. In addition, malaria parasites induce the formation of membranous compartments – the parasitophorous vacuole, the tubovesicular network, the Maurer’s clefts and small vesicles – within the infected erythrocyte, a cell that is normally devoid of internal membranes. After infection, changes also occur in the composition and asymmetry of the erythrocyte plasma membrane. Although many aspects of the mechanism of export of parasite proteins have become clear, the mechanism by which these membranous compartments are formed and expanded is almost entirely unknown. To determine whether parasite-derived phospholipids play a part in these processes, we applied a metabolic labelling technique that allows phosphatidylcholine to be labelled with a fluorophore. As the host erythrocyte cannot synthesize phospholipids, within infected erythrocytes, only parasite-derived phosphatidylcholine will be labelled with this technique. The results revealed that phosphatidylcholine produced by the parasite is distributed throughout the infected erythrocyte, including the TVN and the erythrocyte plasma membrane, but not Maurer’s clefts. Interestingly, labelled phospholipids were also detected in the erythrocyte plasma membrane very soon after invasion of the parasites, indicating that the parasite may add phospholipids to the host erythrocyte during invasion.

## INTRODUCTION

The symptoms of malaria are the result of the infection of erythrocytes by the malaria parasite. Successful infection requires modifications of the host erythrocyte that result in various changes in the host erythrocyte – these allow the parasite to evade the host immune system and elimination in the spleen and increase uptake of nutrients. Many of the best-known modifications (such as those leading to changes in the deformability of the host erythrocyte, its adhesive properties and its ability to take up nutrients) are the result of the export of proteins from the parasite to the host erythrocyte (Maier et al., 2008, 2009; Spillman et al., 2015). Although many questions about the mechanism of protein export remain, several key steps are now well characterised (Beck and Ho, 2021; Garten and Beck, 2021).

In addition to changes in the protein content of the host erythrocyte, the parasite induces the formation of membrane-bound compartments in the erythrocyte –the exomembrane system (Fig 1A) – that includes the Maurer’s clefts, small membranous compartments that are present underneath the host erythrocyte plasma membrane; the parasitophorous vacuole membrane (PVM), which surrounds the parasite and separates it from the host erythrocyte; the tubovesicular network (TVN), a little-studied extension of the parasitophorous vacuole membrane that protrudes into the host erythrocyte cytosol, frequently in a lollipop shape; and small vesicles that are often detected near the Maurer’s clefts (Sherling and van Ooij, 2016). The origin of the membranes that form these structures is in most cases not known. Investigation of the invasion process has indicated that the nascent PVM that surrounds the parasite during and immediately after invasion is derived of host erythrocyte membrane (Geoghegan et al., 2021), but the origin of the phospholipids required for the formation and expansion of the other internal membranes is unclear. The parasite can take up exogenous phospholipids (Moll et al., 1988; Haldar et al., 1989; Fraser et al., 2021; Patel et al., 2022), which the parasites may use to build these compartments, but the parasite also grows in the absence of exogenous phospholipids when the medium is supplemented with fatty acids (Divo et al., 1985; Mitamura et al., 2000; Asahi et al., 2005; Mi-Ichi et al., 2006, 2007; Asahi, 2009). Hence, the parasite appears able to fulfil its phospholipid requirement solely with phospholipids that it synthesizes itself (Vial et al., 1990). In accordance with this, the parasite encodes all the enzymes required for the *de novo* synthesis of phospholipids (Déchamps et al., 2010a). This includes the enzyme for the synthesis of phosphatidylcholine (choline/ethanolamine-phosphotransferase (CEPT)), which is essential and is located in the endoplasmic reticulum of the parasite, as determined in the rodent malaria parasite *Plasmodium berghei* (Déchamps et al., 2010b)

**Figure 1:**
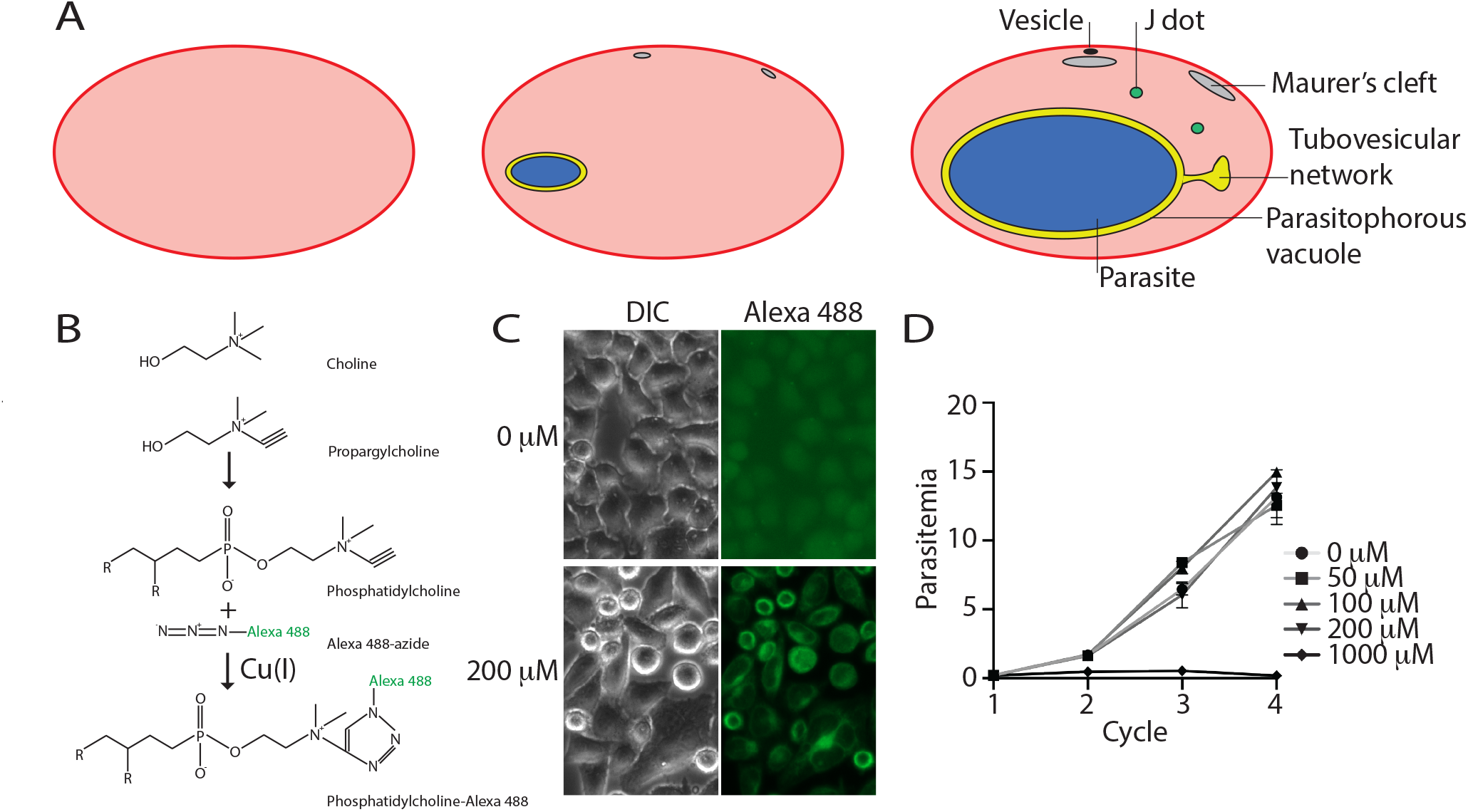
A) Erythrocytes before and after infection with a *Plasmodium falciparum* parasite. The uninfected erythrocyte (left) is devoid of internal organelles, whereas an erythrocyte infected with a young parasite (center) contains multiple internal organelles. An erythrocyte infected with a late-stage parasite (right) contains further organelles. The origins of the lipids that forms these membranes has not been identified. B) Outline of the metabolic incorporation of propargylcholine into phosphatidylcholine and subsequent labelling with Alexa 488-azide using click chemistry. Propargylcholine can be metabolically incorporated into phosphatidylcholine, which can subsequently be conjugated to Alexa 488-azide using Cu(I) as catalyst, forming fluorescent phosphatidylcholine-Alexa 488. C) Labelling of HeLa cells with Alexa 488-azide that had been cultured in the presence of the indicated concentration of propargylcholine for 48 h. D). Growth of *P. falciparum* parasites in the presence of the indicated concentration of propargylcholine over four growth cycles. The experiment was performed in triplicate. Shown is one of two biological replicates.

In addition to the formation of membranous structures, the parasite induces changes in the composition and architecture of the erythrocyte plasma membrane. The distinct phospholipid asymmetry in the erythrocyte plasma membrane is altered in infected erythrocytes, with the level of phosphatidylethanolamine (PE) and phosphatidylcholine (PS) increasing in the outer leaflet (Joshi et al., 1987; Schwartz et al., 1987; van der Schaft et al., 1987; Hsiao et al., 1991; Maguire et al., 1991; Fraser et al., 2021). The phospholipid asymmetry in the plasma membrane of the erythrocyte is maintained by host erythrocyte flippase, floppase and scramblase activity (Hankins et al., 2015). However, after infection by a malaria parasite, the activity of flippase and scramblase changes, leading to the changes in phospholipid level in the different membrane leaflets (Fraser et al., 2021). The fluidity of the membrane increases in the infected erythrocytes, which correlates with the age of the parasites (Taraschi et al., 1986; Joshi et al., 1987). In addition, the level of cholesterol and of sphingomyelin in the erythrocyte plasma membrane decrease as the parasite matures (Maguire and Sherman, 1990; Maier and van Ooij, 2022). Furthermore, the acyl chain composition of the plasma membrane phospholipids is altered in infected erythrocytes compared to uninfected erythrocytes; palmitic acid (16:0) and oleic acid (18:1) become more prevalent in the plasma membrane of infected erythrocytes (Simões et al., 1990, 1992; Hsiao et al., 1991; Wunderlich et al., 1991). Interestingly, palmitic acid and oleic acid are the most prevalent fatty acids in phospholipids in the parasite, which has led to speculation that the parasite either modifies the phospholipids in the erythrocyte plasma membrane or transports phospholipids there (Hsiao et al., 1991; Wunderlich et al., 1991; Simões et al., 1993).

Overall, it remains unclear how the internal membranes are formed and expanded in the host erythrocyte and how the parasite alters the composition of the host erythrocyte membrane. To gain insight into the origin of the phospholipids that form the compartments of the exomembrane system and the alterations of the host erythrocyte plasma membrane, we applied a metabolic labelling technique that allows newly synthesized phosphatidylcholine to be labelled using click chemistry. This revealed that parasite-derived phosphatidylcholine is distributed throughout the host erythrocyte but is not present in Maurer’s clefts.

## MATERIALS AND METHODS

### Parasites

All experiments were performed with the *Plasmodium falciparum* strain 3D7 and the *Plasmodium knowlesi* strain A1-H.1 (Moon et al., 2013). The parasites were maintained as described previously (Ressurreição et al., 2020). Briefly, the parasites were maintained at 37°C in RPMI-1640 (Life Technologies) supplemented with 0.5% AlbuMax type II (Gibco), 50 μM hypoxanthine and 2 mM L-glutamine. For *Plasmodium knowlesi* the medium was further supplemented with 10% (v/v) horse serum. *P. falciparum* was incubated in the presence of 5% CO_2_, *P. knowlesi* was incubated in an atmosphere of 90% N_2_, 1% O_2_, 3% CO_2_. The hematocrit of the cultures was 2-3%. Human erythrocytes were obtained from the National Blood Transfusion Service, UK, and from Cambridge Bioscience, UK.

*P. falciparum* parasites were synchronized by isolating late-stage infected erythrocytes on a discontinuous Percoll gradient. These isolated parasites were washed with RPMI-1640 and mixed with fresh erythrocytes to allow invasion. After 2-3 h, the remaining late-stage parasite were removed using a discontinuous Percoll gradient and the resulting ring-stage culture was treated with 5% D-sorbitol to remove any remaining late-stage parasites (Radfar et al., 2009; Childs et al., 2013). To examine the parasites during and after invasion, schizonts were isolated on a Percoll gradient and incubated at 37°C in the presence of 25 nM ML10 to prevent egress. ML10 was then removed by centrifugation of the culture at 800 x g for three min and resuspension of the erythrocytes in warm RPMI-1640 with supplements. The parasites were then added to fresh erythrocytes, allowing for rapid and highly synchronous invasion (Ressurreição et al., 2020, 2022). To block invasion, cytochalasin D was added to the invading parasites at a concentration of 2 μM.

Growth of parasites was determined using a standard growth assay over four growth cycles. Synchronized ring-stage parasites cultures were adjusted to a parasitemia of 0.1% and maintained in a 24-well dish. Propargylcholine was added at the concentration indicated from a 100 mM aqueous stock solution. A 50 μl aliquot of the culture was removed every 48 h and mixed with 50 μl of 2x fixative solution (8% paraformaldehyde, 0.2% glutaraldehyde and 2x SYBR Green in PBS). Fixed parasites were stored at 4°C until all samples were obtained. Starting from the third cycle, the medium was changed daily (the presence of propargylcholine was maintained throughout). When all samples had been obtained, the parasites were pelleted at ∼2,000 x g for one minute and the fixative was replaced with 1 ml PBS. A 200 μl aliquot was transferred to a 96-well plate and the parasitemia was determined using a Thermo Attune NxT cytometer equipped with a Cytkick Max autosampler. Laser settings used for detection of the erythrocytes and parasites were as follows: forward scatter 125 V, side scatter 350 V, blue laser (BL1) 530:30 280V. The experiment was set up in triplicate, with two biological replicates.

Plasmid pS156 was introduced into 3D7 parasites using transfection of late-stage schizonts as described previously (Collins et al., 2013), followed by selection with 300 μg/ml G418. The plasmid pS156 was a kind gift of Sarah Tarr and Andrew Osborne (University College London). It was produced by replacing the gene encoding the Rex3-GFP fusion and *bsd* in the plasmid PFI1755c 1–61:GFP (Tarr et al., 2013) with a gene encoding SBP1-mCherry and *neo*, respectively.

### Phospholipid labelling

Phosphatidylcholine was labelled using propargylcholine metabolic labelling and subsequent click chemistry as described previously (Salic et al., 2008; Jao et al., 2009). Propargylcholine was added directly to the culture medium of parasites maintained in a 24-well dish; unless otherwise stated, the concentration used was 200 μM, diluted from a 100 mM aqueous stock solution. Parasites were harvested and washed with RPMI-1640 before resuspension in 1 ml fixative (4% paraformaldehyde, 0.1% glutaraldehyde in PBS) and incubation at room temperature for one hour on a rotating wheel. It was noted that increasing the ratio of fixative to parasites improved the subsequent labelling. Parasites were pelleted (all spins in the procedure were at 2,000 x g for one minute) and the fixative was replaced with 1 ml PBS. To label phosphatidylcholine containing propargylcholine with Alexa 488-azide, the parasites were pelleted, resuspended in 500 µl of freshly prepared labelling mix (40 mM Tris, pH 8.5, 2 mM CuSO_4_, 10 µM Alexa 488-azide, 2 µg/ml wheat germ agglutinin (WGA)-Alexa 647), 2 µg/ml Hoechst 33342 and 100 mM ascorbic acid (the ascorbic acid was always added last)) and incubated at room temperature for 30 min on a rotating wheel. The parasites were then washed six-eight times with 1 ml PBS. Fixed and labelled parasites were stored in PBS at 4°C. To prepare the parasites for microscopy, 1.5 µl of parasite pellet was mixed with 1.5 µl Vectashield mounting medium on a microscope slide. The mixture was covered with a coverslip and sealed with nail polish.

### HeLa cell culture

HeLa cells were maintained in RPMI-1640 supplemented with 5% fetal bovine serum in an atmosphere of 5% CO_2_. Cells were maintained using periodic trypsinization and dilution. To label HeLa cells, the cells were seeded on round, acid-treated coverglasses in a 24-well dish and incubated in the presence of 200 μM propargylcholine for approximately 48 h, starting when the cells were ∼40% confluent. The cells were then fixed and labelled with Alexa 488-azide similar to the fixation and labelling of erythrocytes described above, as described previously (Salic et al., 2008; Jao et al., 2009).

### Microscopy

Samples were imaged on a Nikon Ti-E inverted microscope with a Hamamatsu ORCA-Flash 4.0 Camera and Piezo stage driven by NIS elements version 5.3 software using the following excitation wavelengths for fluorescence image acquisition: 365 nm (Hoechst), 470 nm (Alexa 488), 580nm (mCherry) and 635nm (WGA). The images were deconvolved using Nikon NIS software and brightness and contrast were set using FIJI. The images were cropped using FIJI and the file size was changed using Photoshop, figures were produced using Illustrator. For confocal microscopy, samples were imaged on a Zeiss LSM 880 confocal microscope driven by Zen Black version 2.3 software. The emission wavelengths used were 462 nm (Hoechst), 548 nm (Alexa 488) and 637 nm (mCherry). Frame time was 11.3246 seconds. Brightness, contrast and crop size were set in FIJI, the file size was changed using Photoshop and figures were produced using Illustrator.

## RESULTS

### Malaria parasites replicate in the presence of propargylcholine

To determine whether the parasite distributes its own phospholipids within the host erythrocyte, we aimed to localize parasite-derived phosphatidylcholine by metabolically labelling phosphatidylcholine using propargylcholine. Propargylcholine is a derivative of choline that is modified with an alkyne group (Fig 1B); this can be incorporated into phosphatidylcholine and subsequently allows a reporter molecule containing an azide group to be covalently linked to it using click chemistry (Fig 1B). Labelling eukaryotic cells with propargylcholine is a well-established technique to investigate the intracellular distribution of phosphatidylcholine (Salic et al., 2008; Jao et al., 2009) and has previously been applied to the investigation of the origin of lipids obtained by malaria parasites during the liver stage (Itoe et al., 2014). We initially tested whether HeLa cells could incorporate propargylcholine and phosphatidylcholine could be visualized using Alexa 488-azide by culturing these cells in the presence or absence of propargylcholine and subsequently labelling these cells using the established protocol (Salic et al., 2008; Jao et al., 2009). This revealed that HeLa cells that had been cultured in the presence of propargylcholine became brightly labelled with Alexa 488-azide, with a staining pattern very similar to that previously detected (Jao et al., 2009) (Fig 1C).

To determine whether it would be possible to label malaria parasites with propargylcholine, we investigated the effect of propargylcholine on the growth of *Plasmodium falciparum* parasites. In the presence of propargylcholine at concentrations of 200 µM or below, the parasites replicated at a rate similar to that of untreated parasites (Fig 1D). Subsequent experiments were therefore carried out using 200 µM propargylcholine unless otherwise stated.

### Parasite-derived phospholipids are transported to the erythrocyte plasma membrane

We next determined whether phospholipids can be visualized in parasites incubated in the presence of propargylcholine. As erythrocytes do not produce phospholipids (Marks et al., 1960), they will not incorporate propargylcholine into phospholipids and hence in erythrocytes infected with malaria parasites, any labelled phosphatidylcholine will have originated in the parasite. Phosphatidylcholine is the most abundant phospholipid in the parasite and makes up approximately 40-50% of the phospholipids produced by the parasite (van der Schaft et al., 1987; Wunderlich et al., 1991), hence, a large fraction of the parasite-derived phospholipids will contain propargylcholine. To label phosphatidylcholine, *P. falciparum* cultures were propagated either in the presence of propargylcholine or without added propargylcholine for 72 h (one-and-a-half cycles), fixed and then labelled with Alexa 488-azide. The plasma membrane of the erythrocytes was also stained with wheat germ agglutinin (WGA) fused to Alexa Fluor 647 to provide a clear outline of the host erythrocyte. Infected erythrocytes in the culture maintained without propargylcholine were not fluorescent after fixation and staining, indicating that Alexa 488-azide labelling of erythrocytes or parasites requires propargylcholine (Fig 2A). In contrast, parasites maintained in the presence of propargylcholine for 72 h were fluorescently labelled (Figs 2B and 2C). This labelling was detected within the parasites, as would be expected in growing parasites that are actively synthesizing phospholipids – late-stage parasites, especially segregated schizonts, were often very brightly labelled (Fig 2C) as demand for newly synthesized phospholipids increases during this stage, owing to the formation of the plasma membrane for the invaginating parasite and the inner membrane complex (Vial et al., 1982). Furthermore, fluorescence was also detected in the erythrocyte plasma membrane, indicating that parasite phospholipids are transported to the host (Figs 2B and 2C). In contrast, uninfected erythrocytes, which are not capable of phospholipid biosynthesis, were not fluorescent, indicating that active phospholipid metabolism is required for the incorporation of propargylcholine and subsequent labelling with the fluorophore.

**Figure 2:**
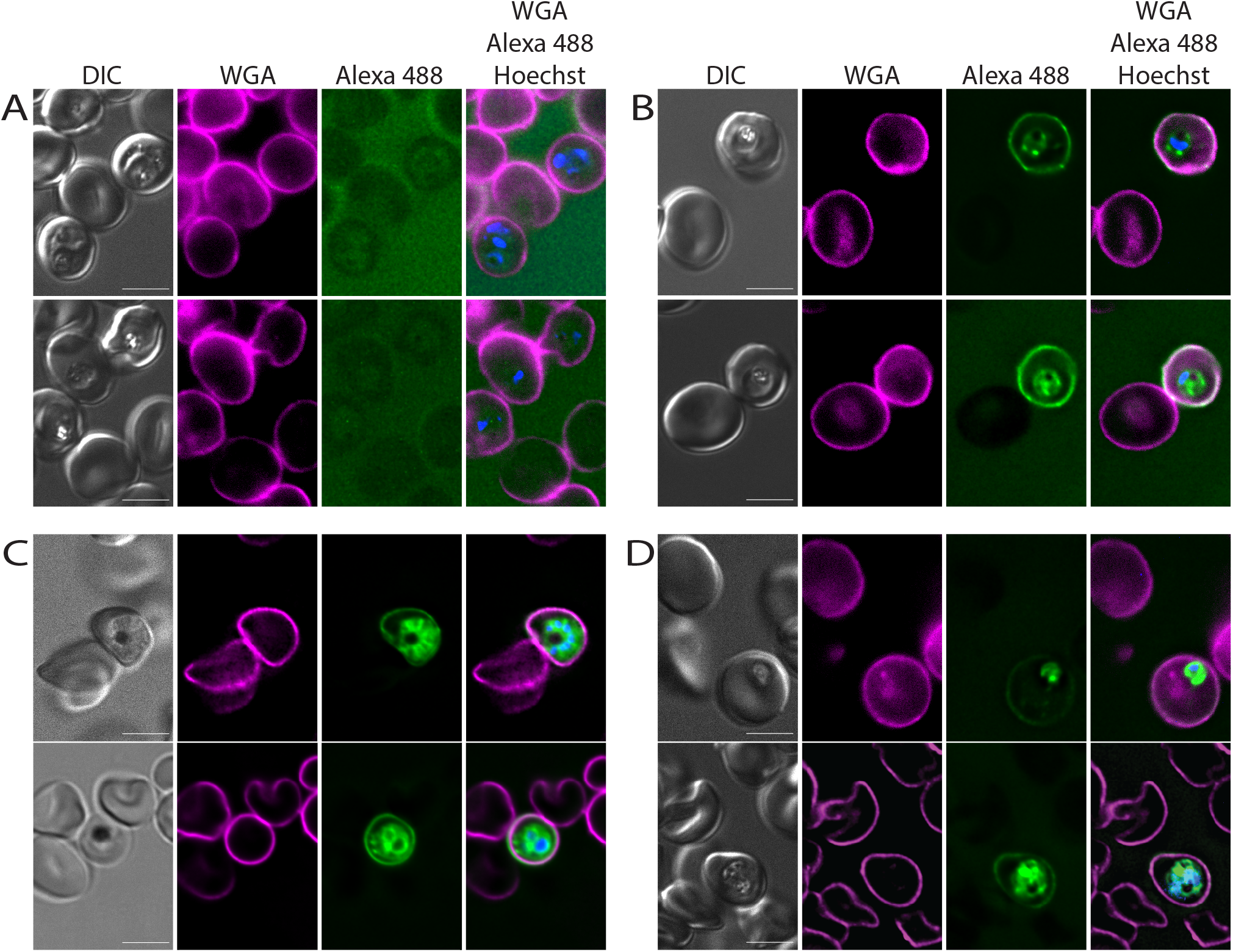
Metabolic labelling of erythrocytes infected with *Plasmodium* parasites with propargylcholine and stained with Alexa 488-azide. A) Erythrocytes infected with *Plasmodium falciparum* were incubated without propargylcholine for 72 h and subsequently fixed and stained with Alexa 488-azide, wheat germ agglutinin (WGA) and Hoechst 33342. B, C) Erythrocytes infected with *Plasmodium falciparum* were incubated in the presence of 200 µM propargylcholine for 72 h and subsequently fixed and stained with Alexa 488-azide, wheat germ agglutinin (WGA) and Hoechst 33342. Shown are trophozoites (B) and schizonts (C), clearly displaying labelling within the parasite and the erythrocyte membrane. No fluorescence was detected in uninfected erythrocytes. D) Erythrocytes infected with *Plasmodium knowlesi* were incubated in the presence of 200 µM propargylcholine for 36 h and subsequently fixed and stained with Alexa 488-azide, wheat germ agglutinin (WGA) and Hoechst 33342. Note the distinct staining of the parasite and erythrocyte plasma membrane in infected erythrocytes and the absence of staining of uninfected erythrocytes. Scale bar: 5 µm.

Decreasing the concentration of propargylcholine in the medium led to lower levels of fluorescence after fixation and labelling (Supp Fig 1). Similarly, when synchronized parasites were incubated in the presence of propargylcholine for only three hours during the late stage of the erythrocytic cycle, only a very low level of fluorescence was detected in the infected erythrocytes as compared to parasites that had been labelled for nearly the entire erythrocytic cycle (Supp Fig 2), indicating that unincorporated propargylcholine was removed either in the initial washing step or subsequent steps of the procedure and the observed fluorescence does not represent propargylcholine that remained in the infected erythrocyte. No labelling of the host erythrocyte membrane was detected in the infected erythrocytes that had been labelled for only a short period.

To determine whether the export of phospholipids is common among *Plasmodium* spp., we repeated the labelling with erythrocytes infected with *Plasmodium knowlesi*. Similar to *P. falciparum*, fluorescence was detected in *P. knowlesi* parasites as well as the infected erythrocyte plasma membrane, but not in uninfected erythrocytes, indicating that the transport of parasite-derived phospholipids to the host erythrocyte plasma membrane is a property shared among *Plasmodium* spp. (Figure 2D).

### Transport of phospholipids to the erythrocyte plasma membrane can be detected from the trophozoite stage

As the preceding experiments were performed over one-and-a-half cycles, it could not be determined at which time after invasion phosphatidylcholine transport of parasite phospholipids to the erythrocyte initiates. Therefore, we added propargylcholine to a culture of synchronized parasites immediately after the parasites had invaded the erythrocytes. Labelling in the erythrocyte membrane was first detected at the trophozoite stage, approximately 34 h after invasion, indicating that phospholipid transfer to the host erythrocyte has initiated at this stage (Fig 3A). Phospholipid transfer possibly initiates earlier, but the level of phospholipid remains below the level of detection during that time; labelling of the erythrocyte plasma membrane was not robustly detected 27 h after invasion (Fig 3A). The level of observed fluorescence detected gradually increased as the parasite matured, with labelling in the plasma membrane of erythrocytes containing schizonts brightest. These results are consistent with the findings that the alteration in the phospholipid content of the host erythrocyte plasma membrane changes most noticeably during the trophozoite stage (Taraschi et al., 1986).

**Figure 3:**
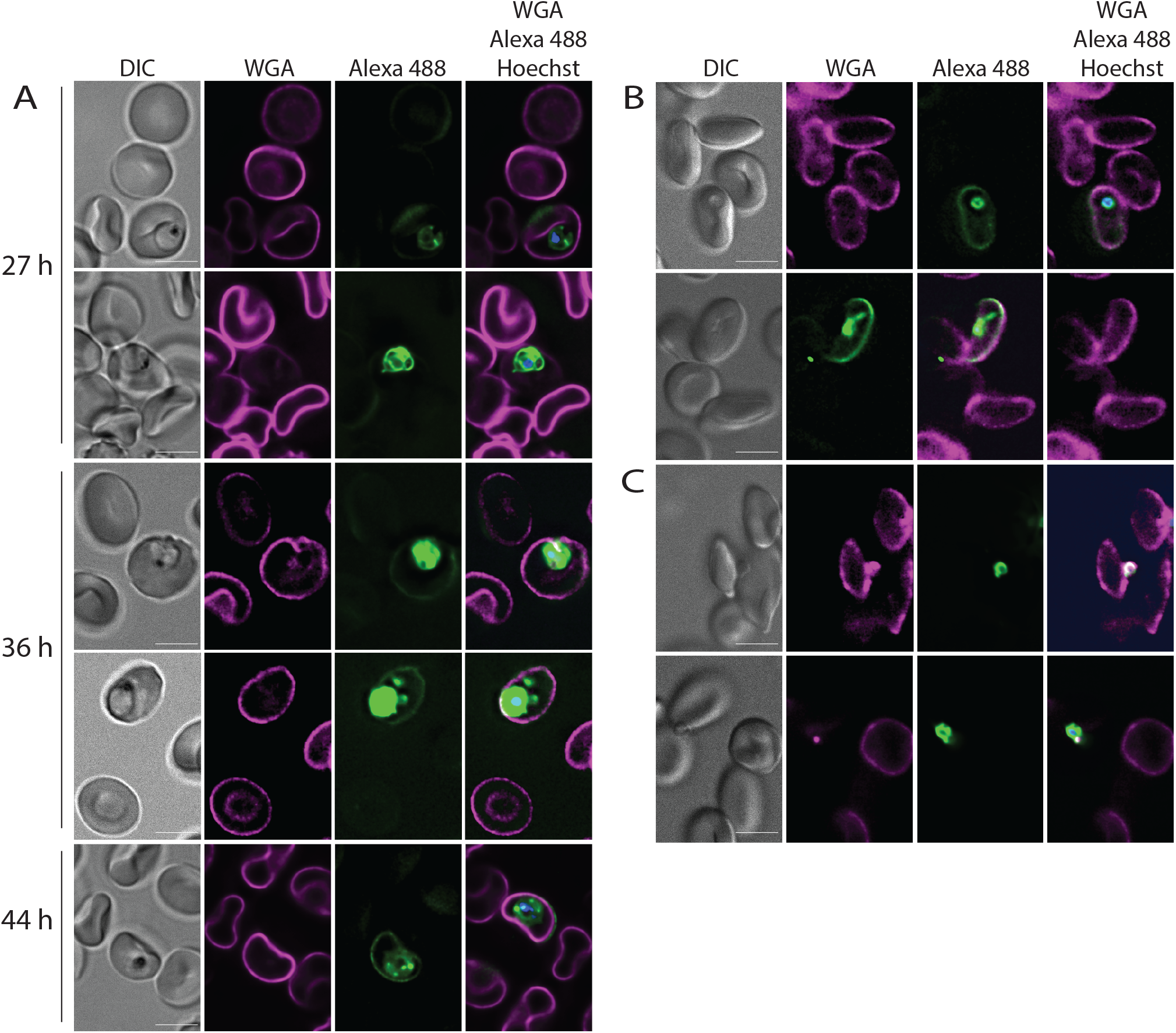
Timing of phospholipid transfer from parasite to the erythrocyte plasma membrane. A) Metabolic labelling of *Plasmodium falciparum* parasites was initiated immediately after invasion using a highly synchronized parasite culture. Samples were removed at the indicated time and fixed. B) Phospholipid transfer to the host erythrocyte after invasion. Schizonts that had been labelled with propargylcholine for 72 h and prevented from egressing by the addition of ML10 were washed, thereby removing the ML10, then allowed to egress and form new rings in the absence of propargylcholine. D) Inhibition of invasion with cytochalasin D prevents transfer of phospholipids from the parasite to the host cell. Parasites were prepared as in C) except that cytochalasin D was added to the culture of egressing parasites. Note the absence of staining of the erythrocyte plasma membrane. Scale bar: 5 µm.

### Phospholipids are transferred from the parasite to the host erythrocyte during or shortly after invasion

The origin of the membrane of the nascent PVM has been a subject of debate (Sherling and van Ooij, 2016), although recent results support a model in which the erythrocyte plasma membrane constitutes the main source of phospholipids (Geoghegan et al., 2021). However, the observed labelling of the host erythrocyte membrane could in part be the result of exchange of phospholipids during or immediately following the invasion process. As the labelling with propargylcholine allows the transfer of parasite phosphatidylcholine to the host erythrocyte to be investigated, we determined the distribution of parasite phosphatidylcholine around the time of invasion. We labelled synchronized parasites with propargylcholine for 72 h and then blocked egress with the cGMP-dependent kinase (PKG) inhibitor ML10 so that the timing of invasion of the labelled parasites could be carefully controlled (Baker et al., 2017; Ressurreição et al., 2020, 2022). These parasites were isolated on a discontinuous Percoll gradient and then allowed to egress in the absence of ML10 and propargylcholine and invade fresh erythrocytes for one hour. This revealed that the plasma membrane of the infected erythrocyte could be fluorescently labelled almost immediately after invasion (Fig 3C, Supp Fig 3), indicating that transfer of phospholipids from the parasite to the host erythrocyte occurs during or very soon after invasion.

To narrow down the stage of the invasion process at which the transfer of phospholipids occurs, we blocked invasion with the actin polymerization inhibitor cytochalasin D (CD). In the presence of CD the parasite binds to and reorients on the erythrocyte surface, forming a tight junction, but does not invade the host erythrocyte (Miller et al., 1979; Weiss et al., 2015); CD therefore inhibits one of the last steps prior to the initiation of invasion. In parasite cultures treated with CD, the parasites, but not the host erythrocytes, were brightly labelled with Alexa 488-azide, indicating that no phospholipid transfer had taken place (Fig 3C). Hence, the transfer of phospholipids from the parasite to the erythrocyte plasma membrane occurs after the parasite has initiated invasion of the host erythrocyte.

### The tubovesicular network but not Maurer’s clefts contains parasite-derived phosphatidylcholine

The parasite induces the formation of multiple internal membranes (Fig 1A). To determine whether these contain parasite-derived phospholipids, examination of the infected erythrocytes revealed that in some cases, loops protruding from the parasites could be detected. These likely represent the TVN (Fig 4A) (Lauer et al., 1997; Wickham et al., 2001; Adisa et al., 2003; Charnaud et al., 2018). This labelling indicates that this structure is also produced, at least in part, with phospholipids produced by the parasite. As the TVN is an extension of the PVM (Elford et al., 1995; Wickham et al., 2001; Adisa et al., 2003), it is likely that the PVM is also composed of parasite-derived phospholipids.

**Figure 4:**
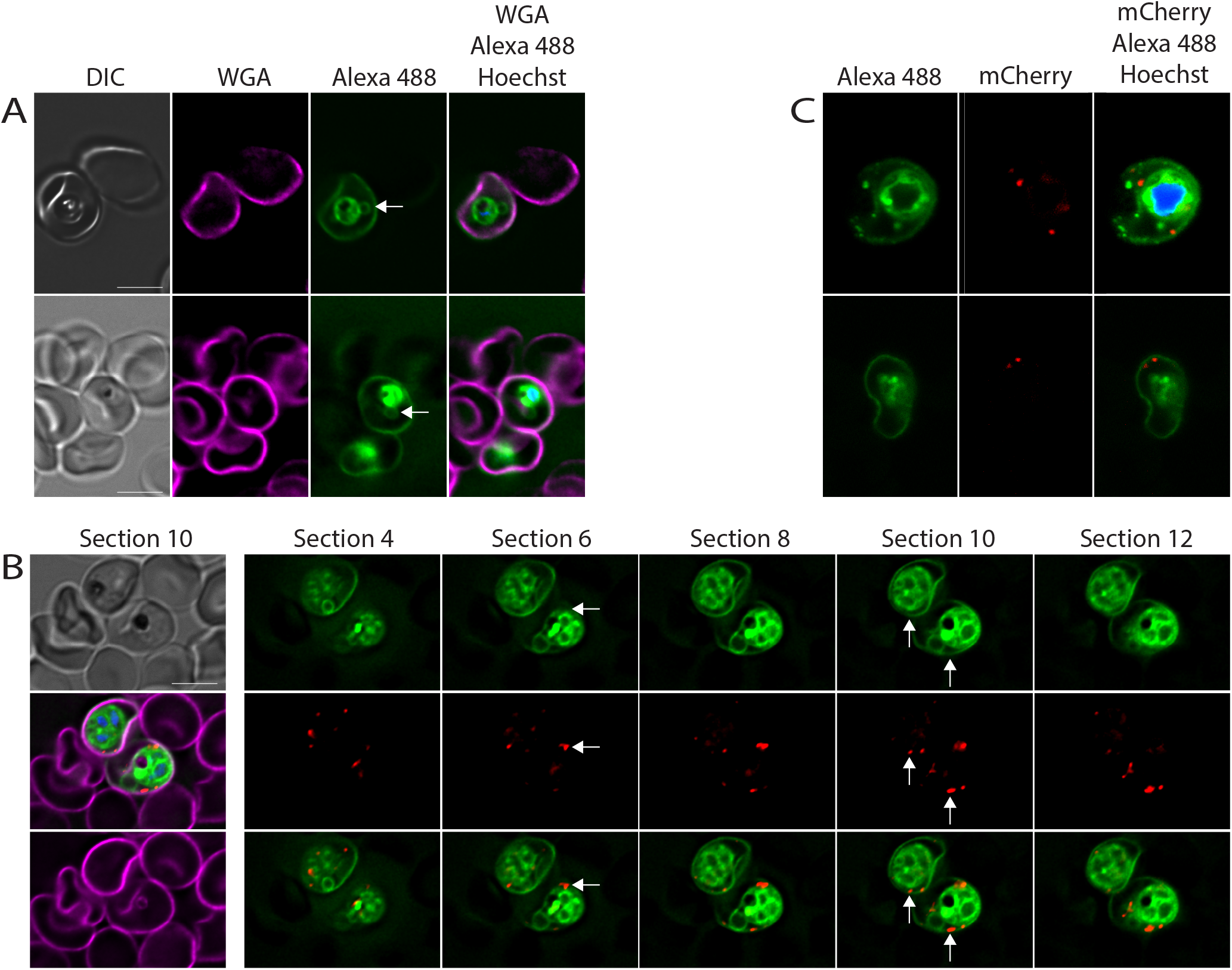
Labelling of the exomembrane system. A) Labelled parasites displaying a stained tubovesicular network (arrow). B) Maurer’s clefts are not labelled with Alexa 488-azide. *Plasmodium falciparum* parasites that produce a fusion of the Maurer’s cleft marker SBP1 with mCherry were metabolically labelled with propargylcholine for 72 h, fixed and stained with Alexa 488-azide. The left-hand column shows section 10 of the deconvoluted z-series, the other columns show the Alexa 488-azide labelling (top), mCherry labelling (middle) and the merge of the Alexa 488-azide and mCherry staining (bottom) of the indicated deconvolved sections. Arrows indicate select Maurer’s clefts that do not overlap with Alexa 488-azide staining. C) Confocal imaging of labelled parasites expressing SBP1-mCherry. Note the absence of colocalization of Alexa 488-azide and mCherry labelling in B) and C). For a 3D view of the parasites shown in C), see supplementary movies 1 and 2. Scale bar: 5 µm.

In the course of our investigation, we did not detect labelling that may represent Maurer’s clefts. To determine whether Maurer’s clefts contain parasite-derived lipids, we repeated the labelling in parasites that produced the Maurer’s cleft protein SBP1 fused to mCherry. As mCherry remains fluorescent after fixation, this allowed visualization of Maurer’s clefts without the need to permeabilize the erythrocyte for antibody staining. Even when the parasites were labelled for one-and-a-half cycles to maximize labelling, we were not able to detect Alexa 488-azide in Maurer’s clefts after staining (Fig 4B). Furthermore, we applied confocal microscopy and 3D reconstruction to look at this potential lack of colocalization in more detail. In this case, too, we were unable to detect staining in the Maurer’s clefts, even though the parasite and the erythrocyte surface were clearly labelled (Fig 4C and supplementary movies 1 and 2). As previous findings had indicated that there is likely to be a difference in the phospholipid content of the PVM and the Maurer’s clefts based on their different permeability after treatment with Streptolysin O (Mundwiler-Pachlatko and Beck, 2013), this finding is perhaps not surprising.

The absence of labelling from the Maurer’s cleft membrane, an internal membrane in the infected erythrocyte, further emphasizes the specificity of the labelling in the other internal and external membranes and underscores that the observed fluorescence reflects the presence of fluorescent phospholipid rather than non-specific retention of propargylcholine or Alexa 488-azide in membranes.

## DISCUSSION

Together, these experiments reveal that phosphatidylcholine produced by malaria parasites is transported from the parasite throughout the infected erythrocyte. Parasite-derived phosphatidylcholine was detected in the TVN (and hence is very likely to be present in the PVM, as the two membranes are physically connected) and the erythrocyte membrane. Transfer of phospholipids to the erythrocyte was detected immediately after invasion and although no transfer could be detected in the ring stage, phospholipid transfer from the parasite to the erythrocyte could be detected at the trophozoite stage. Interestingly, the Maurer’s clefts did not contain any detectable fluorescence after labelling, indicating that these structures are not produced using parasite phosphatidylcholine. The origin of the Maurer’s clefts has remained unclear; early suggestions that they are derived from the PVM (or even joined to the PVM) were contradicted by the finding that the PVM and Maurer’s clefts differ in their sensitivity to streptolysin O (Mundwiler-Pachlatko and Beck, 2013), indicating that their membrane compositions differ. Our results indicate that these compartments may form very soon after invasion of the erythrocyte and not be produced with parasite phospholipids.

The mechanism of the transport of phospholipids through the host erythrocyte remains to be discovered. As phosphatidylcholine is synthesized in the ER of the parasite (Déchamps et al., 2010b), it is transported through the secretory pathway of the parasite to the parasite plasma membrane. From there, it must be transported through the aqueous lumen of the PV to the PVM – this could potentially occur through membrane contact sites between the parasite plasma membrane and the PVM (Garten et al., 2020) – from where the phospholipid has to be transported to the erythrocyte plasma membrane. The mechanism by which this occurs is not obvious. One candidate for a phospholipid transporter is the J dot, which carries cholesterol (Behl et al., 2019) but to date has not been shown to contain phospholipids. Alternatively, phospholipids could be carried by a phospholipid transfer protein (Ressurreição and van Ooij, 2021). However, in contrast to the role of Maurer’s clefts in transport of proteins to the RBC plasma membrane, proteins, it is clear that Maurer’s clefts do not form an intermediate stage for transport of phosphatidylcholine to the surface of the erythrocyte, as no parasite-derived phosphatidylcholine was detected in Maurer’s clefts.

Previous studies have shown that the phospholipid composition in the host erythrocyte plasma membrane changes the biophysical properties and fatty acid composition of the phospholipids of the infected erythrocyte plasma membrane (Taraschi et al., 1986; Simões et al., 1990; Wunderlich et al., 1991). Using the biochemical methods employed, it could not be distinguished whether this was the result of transport of parasite-derived lipids to the membrane or whether reflected modification of host lipids. Our finding that parasite-derived phosphatidylcholine is transported to the host erythrocyte plasma membrane likely indicates that the change in membrane lipids is the result of addition of parasite phospholipids to the erythrocyte plasma membrane. Hence, the modification of the properties of the host erythrocyte are likely to be altered by the parasite not solely by the export of proteins, but also through the export of phospholipids.

## ACKNOWLEDGMENTS

The work in this study was supported by a Career Development Award from the Medical Research Council to CvO (MR/R008485/1) and a Medical Research Council London Intercollegiate Doctoral Training Partnership Studentship to TV. The plasmid pS156 was a kind gift of Sarah Tarr and Andrew Osborne (University College London). The authors acknowledge the facilities and the scientific and technical assistance of the LSHTM Wolfson Cell Biology Facility, with specific thanks to Liz McCarthy.

## LEGENDS

Supplementary Figure 1: Titration of propargylcholine. Cultures of infected erythrocytes were incubated in the presence of the indicated concentration of propargylcholine for 72 h. The erythrocytes were then fixed and labelled as described. Scale bar: 5 µm

Supplementary Figure 2: Short-term labelling of erythrocytes infected with *Plasmodium falciparum*. Cultures of *P. falciparum*-infected erythrocytes were incubated in the presence of propargylcholine starting either 2 h after invasion (A) or 42 h after invasion (B). The culture was fixed and labelled 45 h after invasion. The parasites were imaged under identical conditions and the obtained images were analyzed using FIJI. The Alexa 488-azide staining is shown using a contrast range of 150-20000 and in the case of the parasites labelled from 42 h post-invasion, also using a contrast range of 150-2000 (the latter higher contrast image allows visualisation of weakly fluorescent cellular structures). Note the significantly lower brightness of the parasites labelled for only 3 h.

Supplementary Figure 3: Analysis of membrane staining in erythrocytes containing recently invaded parasites. Brightness of erythrocyte membrane in deconvolved images of infected and uninfected erythrocytes was analyzed using the line intensity tool of the Nikon NIS software. Shown in each panel are fluorescence intensity plots of an infected erythrocyte (line 1) and an uninfected erythrocyte (line 2). Shown below is the intensity of the green signal along the line shown in the panel. Scale bar: 5 µm.

Supplementary movie 1: 3D reconstruction of the infected erythrocyte shown in the top panel of Figure 4C.

Supplementary movie 2: 3D reconstruction of the infected erythrocyte shown in the bottom panel of Figure 4C.

