## Supplementary figures and images for "Distribution of malaria parasite-derived phosphatidylcholine in the infected erythrocyte"

### Supplementary figures

Supplementary Figure 1

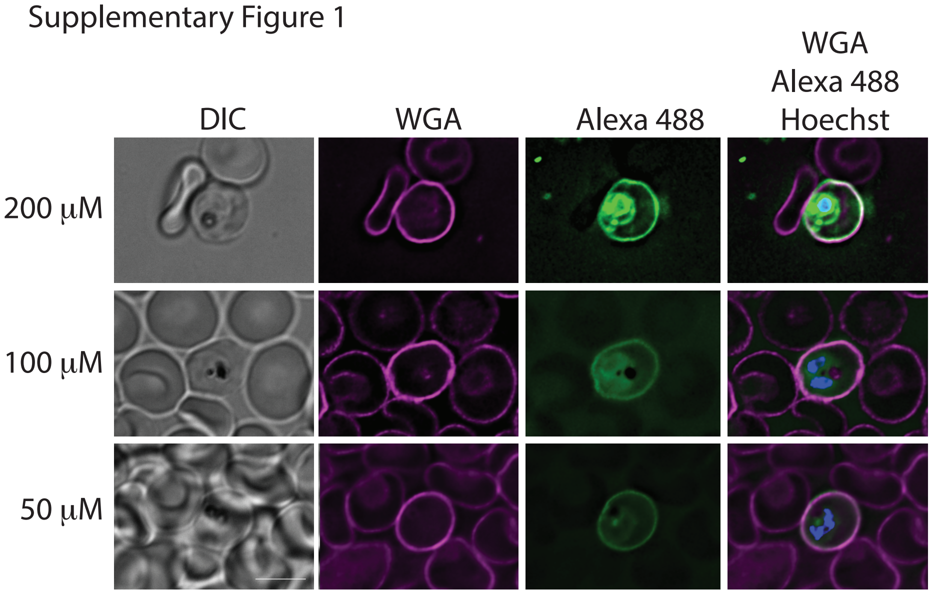
